# Common modes of ERK induction resolve into context specific signalling via a combination of signal duration and cell type specific transcriptional repression

**DOI:** 10.1101/2024.02.29.582771

**Authors:** Marta Perera, Joshua M Brickman

## Abstract

Fibroblast Growth Factor (FGF) signalling via ERK exerts diverse roles in development and disease. In mammalian preimplantation embryos and naïve pluripotent stem cells ERK promotes differentiation, whereas in primed pluripotent states closer to somatic differentiation ERK sustains self-renewal. How can the same pathway produce different outcomes in two related cell types? To explore context-dependent ERK signalling in embryonic development and differentiation we generated cell and mouse lines that allow for tissue- and time-specific cell intrinsic ERK activation. Using these tools, we find variations in response to ERK are mostly mediated by repression of transcriptional targets that occur in tandem with reductions in chromatin accessibility at regulatory regions. Furthermore, immediate early ERK responses are largely shared by different cell types, but become refined producing cell-specific programs as increasing durations of signalling interface with cell specific gene regulatory networks. Induction in naïve pluripotency is accompanied by chromatin changes, whereas in later stages it is not, suggesting that chromatin context doesn’t shape signalling response. Altogether, our data suggest that cell-type specific responses to ERK signalling exploit the same immediate early response, but then sculpt it to specific lineages via repression of distinct cellular programs and downstream indirect stimulation of available enhancer networks.

## Introduction

Cell fate decisions during embryonic development are driven by a limited number of extracellular signals that operate at the precise time and place to activate distinct lineage-specific programs. Given there is a finite number of signals but thousands of different cell types, it becomes necessary to repurpose these signals to produce cell type-specific outcomes. Fibroblast Growth Factor (FGF) signalling *via* Extracellular signal-Regulated Kinase (ERK) is a prime example of this problem, exploiting the same set of kinases to produce diverse responses in multiple tissues during embryonic development (Corson et al., 2003; Dorey and Amaya, 2010). FGF/ERK plays a fundamental role in coordinating cell fate choice and is one of the most frequent pathways mutated in cancer (Brewer et al., 2016; Dorey and Amaya, 2010; Ornitz and Itoh, 2015).

Early development in mammals is concerned with the segregation of the embryonic Epiblast from the extra-embryonic lineages required for post implantation development, the Trophectoderm (TE) and the Primitive Endoderm (PrE); and FGF/ERK is required to make both. First, it is important for the segregation of outside cells into the TE from the inner cells into the Inner Cell Mass (ICM) (Lu et al., 2008; Nichols et al., 1998; Nichols et al., 2009). Then, just before implantation at embryonic day (E)3.5, FGF/ERK is necessary to drive segregation of PrE from the Epiblast (Chazaud et al., 2006; Krawchuk et al., 2013; Yamanaka et al., 2010). This segregation is initiated by Epiblast cells secreting FGF4, which activates ERK in a neighbouring cell, inducing PrE identity (Azami et al., 2019; Chazaud et al., 2006; Frankenberg et al., 2011; Kang et al., 2017; Molotkov et al., 2017). As a result, culture of pre-segregation mouse embryos with either FGF4 or an FGF/ERK signalling inhibitor produces an ICM composed entirely of PrE or Epiblast cells, respectively (Nichols et al., 2009; Schrode et al., 2014; Yamanaka et al., 2010).

Based on the capacity of ERK inhibition to block PrE differentiation, mouse Embryonic Stem Cells (mESCs) can be efficiently derived by the *ex vivo* expansion of peri-implantation Epiblast in the presence of an FGF/ERK signalling inhibitor (Ying et al., 2008). This supports their expansion as cells with the competence to differentiate into all lineages, a property termed pluripotency. As the gene expression state in these cells resembles the earliest stage of embryonic development, they are called naïve (Nichols and Smith, 2009). Stimulation of ERK induces naïve cells to exit naïve pluripotency, supressing the expression of factors known to support pluripotency at the same time as activating a differentiation program by acting directly on enhancer activity (Hamilton et al., 2019). However, the activity of ERK rapidly changes at implantation. Post-implantation Epiblast cells can also be expanded as pluripotent stem cell cultures, but in these cases FGF/ERK is required for their expansion and to block differentiation into somatic lineages (Brons et al., 2007; Tesar et al., 2007). Their gene expression profiles approximate later stages of embryonic development than naïve cells, and therefore are termed primed (Endoh and Niwa, 2022; Morgani et al., 2017; Nichols and Smith, 2009).

FGF/ERK is not the only pathway to promote opposing outcomes in pre-vs post-implantation Epiblast: at the naïve stage WNT/ß-catenin activation (or GSK3 inhibition) blocks differentiation to promote pluripotency, while in primed Epiblast induces differentiation (Sumi et al., 2013; Ying et al., 2008). *In vivo*, the Epiblast progresses out of the naïve state through two morphogenetic changes: first, Epiblast cells become polarized to produce rossette-like structures at E5.0, then a central lumen arises and the cells form a cup-shaped Epiblast at E5.5 (Bedzhov and Zernicka-Goetz, 2014; Shahbazi et al., 2017). The formation of these morphological features has been exploited to identify the temporal hierarchy of WNT and FGF signalling. Naïve mESCs and *ex vivo* cultured embryos can be captured in a rosette-like state in the presence of FGF and WNT inhibition, and then advanced further into Epiblast development upon FGF/ERK release (Neagu et al., 2020). This led to the identification of Rosette Stem Cells (RSCs), analogous to the E5.0 Epiblast (Endoh and Niwa, 2022; Neagu et al., 2020). RSCs can be expanded in the presence of both WNT and FGF inhibitors, and then progressed towards primed pluripotency *via* FGF/ERK activation to generate Epiblast-Like Cells (EpiLCs) (Hayashi et al., 2011). The existence of these defined culture models that mimic the progression through naïve into primed Epiblast development provides an ideal platform to investigate the molecular cues that regulate FGF/ERK signalling context.

In this paper, we have developed mouse and cell culture models that allow time- and cell type-specific ERK activation by exploiting a combination of our intrinsic c-RAF induction model (Hamilton and Brickman, 2014; Hamilton et al., 2019) and Cre-mediated recombination. We have used these models to map responses to ERK signalling as cell states progress from naïve to primed pluripotency. When we examine early signalling responses, we find a striking conservation of inductive events across cell types. However, later signalling responses are more variable between cell types and are mediated by repression of cell type-specific transcription. Finally, we have investigated the mechanism of ERK-mediated context responses by mapping FGF-dependent regulatory regions in each cellular context. In naïve pluripotency, ERK triggers functional chromatin remodelling, while in primed pluripotency, it exploits the existing accessible regions.

## Results

To investigate the cellular responses to ERK in mouse pre- and post-implantation Epiblast, we designed a strategy that allowed us to induce ERK *in vivo* and *in vitro*. We took advantage of a previously published genetic model (Hamilton and Brickman, 2014; Hamilton et al., 2019) which employs a constitutively active c-RAF (*Raf1*) to activate the kinase cascade upstream of ERK (Fig. 1A). The engineered c-RAF kinase is fused to a tamoxifen (4OHT)-specific ligand binding domain of the oestrogen receptor (ER^T2^), allowing for precise temporal activation upon 4OHT supplementation (Fig. 1A). To enable *in vivo* lineage-specific Cre-mediated ERK activation, we introduced a lox-stop-lox cassette upstream of the inducible c-RAF. We furthermore tagged the inducible c-RAF with HA and followed it by a T2A self-cleaving peptide and a H2B-mCherry reporter, allowing us to visualise transgene expression based on either H2B-mCherry fluorescence or HA immunodetection (Fig. 1B). We targeted mESCs and verified ERK induction and mCherry expression (Figs S1A-E) before injecting into morula-stage embryos to produce chimeric mice, which were then crossed to obtain a stable homozygous line. The line was called CRAFR26 (Conditional RAF in Rosa26 locus).

**Figure 1.**
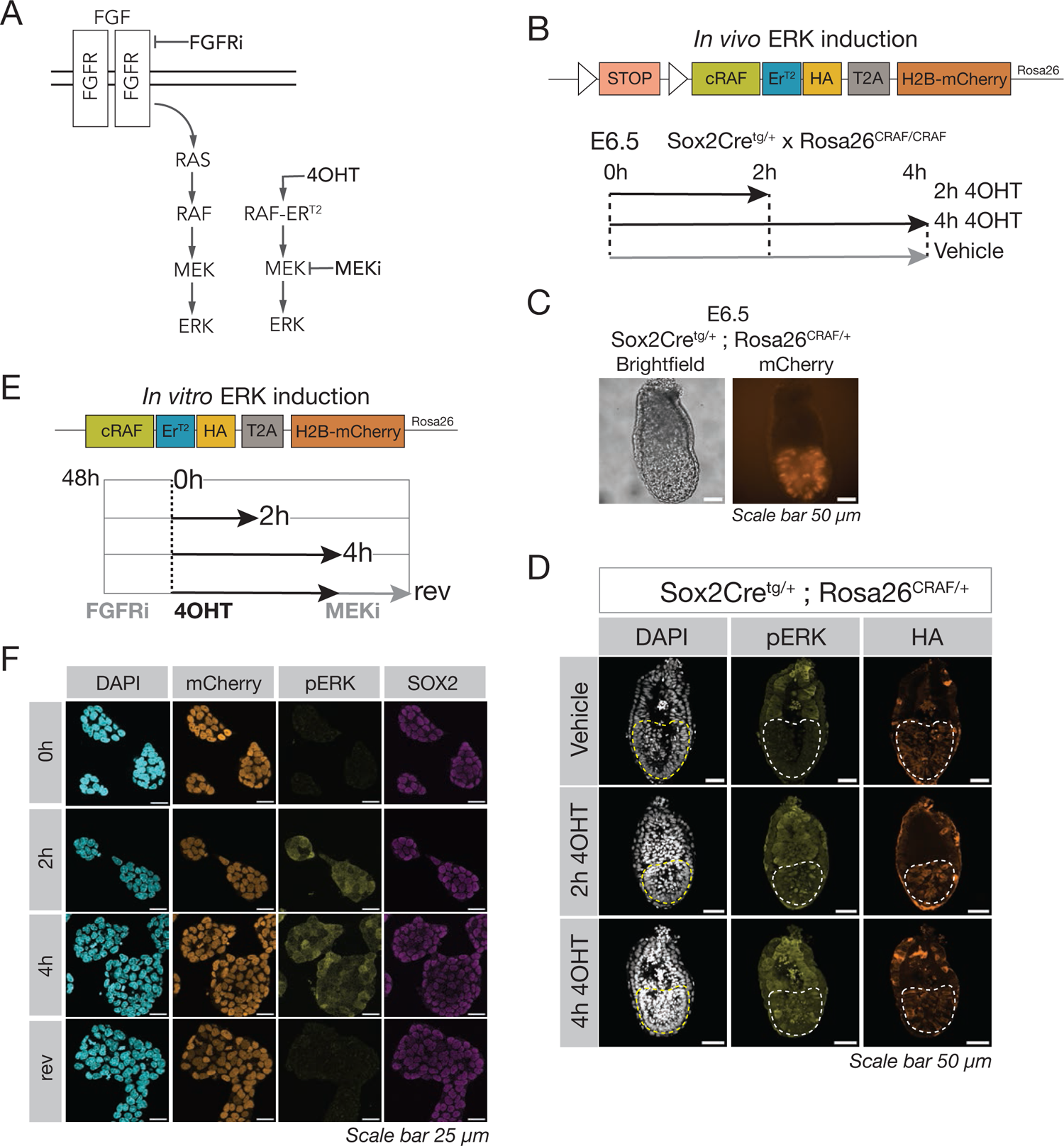
A new mouse and mESC tool to activate ERK in a time- and tissue-specific manner. A. Diagram of the strategy to control ERK activation and inhibition. Culture with FGF Receptor inhibitor (FGFRi) inhibits the endogenous FGF/ERK pathway. The exogenous c-RAF-ER^T2^ is activated with tamoxifen (4OHT). The activation can be rapidly reverted by adding MEK inhibitor (MEKi), upstream of ERK. B. Outline of the CRAFR26 mouse line construct. A lox-stop-lox cassette prevents transcription unless Cre recombinase is expressed. A constitutively active form of the kinase c-RAF produces phosphatase-independent ERK induction. The c-RAF is fused with an ER^T2^, rendering it inactive unless 4OHT is present. An HA-tag allows detection of the construct by Western blot or immunofluorescence. A T2A self-cleaving peptide separates the c-RAF from an H2B-mCherry fluorescent protein that labels all cells that express the construct. To activate the pathway during embryonic development, Sox2Cre heterozygous males were crossed with CRAFR26 homozygous females to obtain Sox2Cre^tg/+^;Rosa26^CRAF/+^ E6.5 embryos. Embryos were then treated with either 250 nM of 4OHT (to activate ERK) or ethanol (Vehicle control) for the indicated times. C. Live imaging of E6.5 Sox2Cre^tg/+^;Rosa26^CRAF/+^ embryos shows expression of the mCherry fluorescent protein in the Epiblast, confirming expression of the CRAFR26 construct restricted to the Epiblast, driven by the Sox2Cre. D. Sox2Cre^tg/+^;Rosa26^CRAF/+^ E6.5 embryos were fixed and stained after the indicated treatment (Vehicle or 4OHT). HA-tag detection shows expression of the construct restricted to the Epiblast driven by the Sox2Cre. pERK staining shows induction of the ERK pathway after 4OHT treatment, and no induction when treated with vehicle. Dashed lines indicate the location of the Epiblast. See quantification and n numbers in Fig. S1G-H. E. The CRAFR26 cell line was transfected with a pCAG-Cre plasmid to remove the lox-stop-lox cassette and allow for constitutive expression of the construct in vitro. The c-RAF kinase is only active upon treatment with 4OHT. CRAFR26 mESCs lines were routinely cultured in 2i/LIF, and 48 hours before the experiment they were plated in FGFRi (see Methods). Induction of the c-RAF kinase was achieved by supplementation of 250 nM of 4OHT to the culture media for the indicated times. Reversion (rev) of the ERK activation was performed by addition of MEKi. Cells were collected at 0h (FGFRi), 2h or 4h of ERK induction, and reversion (4h of induction plus 2h of MEKi). F. CRAFR26 mESCs were fixed at the indicated times of ERK induction and stained for pERK to show ERK induction, and SOX2 to confirm maintenance of mESC pluripotency during the induction. mCherry fluorescence indicates homogeneous expression of the CRAFR26 construct.

To validate the system, we crossed homozygous CRAFR26 females with heterozygous Sox2Cre males (Hayashi et al., 2002), which expresses Cre in the Epiblast lineage. We observed mCherry signal in the Epiblast of E6.5 embryos (Fig. 1C) and induced the c-RAF kinase with 4OHT for 2 or 4 hours. 4OHT-treated embryos produced higher pERK expression in the Epiblast compared to non-treated embryos (Vehicle) (Figs 1D, S1F-H), or non-floxed embryos (Sox2Cre^+/+^) (Figs S1F-H).

To investigate the context-dependent roles of ERK *in vitro*, CRAFR26 mESCs were transiently transfected with Cre, generating a cell line that constitutively expressed *cRAF-HA-T2A-H2B-mCherry* from the *Rosa26* locus (Fig. 1E). The cells show homogeneous induction of pERK after 2 or 4 hours of 4OHT treatment, rapidly reversed by a MEK inhibitor (MEKi, see Methods) (Fig. 1F). Cells were cultured in the presence of FGF receptor inhibitor (FGFRi) for 48 hours before and during the induction, to ensure that the source of ERK signalling comes from the exogenous construct. We detected SOX2 during ERK induction to confirm that cells maintained their ESC identity (Fig. 1F).

### ERK activates an early shared transcriptional response and later cell type-specific programs

To precisely map the transcriptional response to ERK in *in vitro* pluripotent cell types around implantation, we differentiated targeted ESCs into RSCs, which are analogous to the E5.0 Epiblast (Neagu et al., 2020), and EpiLCs, related to the E5.5 Epiblast (Hayashi et al., 2011) (Fig. 2A). We analysed the transcriptome of these cells by bulk RNA-sequencing (RNA-seq) before and after ERK stimulation and reversion with MEKi (Fig. 2A). Principal Component Analysis (PCA) projection shows that the biggest change is driven by developmental time: there is a stepwise progression ESCs-RSC-EpiLCs as cells transition from naïve to primed pluripotency (Fig. 2B).

**Figure 2.**
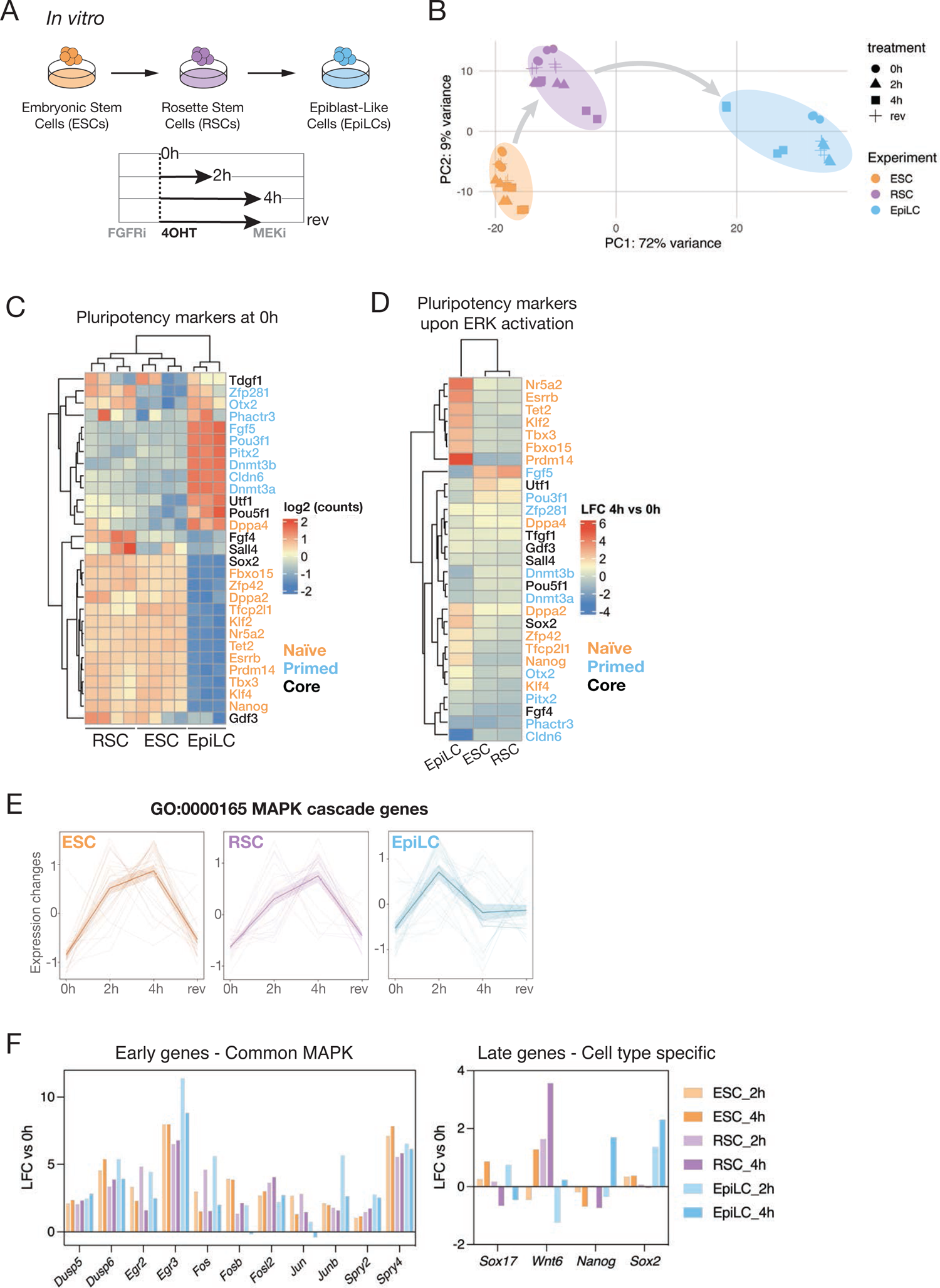
Pluripotent *in vitro* RNA-seq timecourse captures an early universal and a late specific ERK transcriptional response. A. Diagram depicting the *in vitro* cell types collected to investigate the context-dependent responses to ERK in naïve (Embryonic Stem Cells, ESCs), primed (Epiblast-Like Cells, EpiLCs) and intermediate (Rosette Stem Cells, RSCs) pluripotent populations. All samples were cultured in FGFRi for 48 hours before beginning the experiment except for EpiLCs, which were in FGFRi for 2 hours. Then ERK is activated for 2h or 4h by addition of 4OHT. To explore reversion (rev) of ERK target genes, MEKi was added for an additional 2h after the induction. 4 biological replicates were collected for each condition (0h, 2h, 4h and rev). 1 sample of EpiLC 0h was discarded during Quality Control. B. Principal Component Analysis (PCA) of the in vitro samples sequenced. PC1 contains most of the variance (72%) and separates naïve ESCs from primed EpiLCs, with RSCs at the intermediate stage. Symbol represents the treatment course described in Fig 2A. C. Heatmap showing the expression of pluripotency markers (orange for naïve markers, blue for primed and black for shared or core pluripotency) of the in vitro cell types before ERK induction (0h as described in Fig 2A). ESCs and RSCs express mostly naïve markers while EpiLCs show higher expression of primed markers. Each column represents a represents an individual biological replicate. ESCs n=4, RSCs n=4, EpiLCs n=3. The colour scale indicates normalised counts in log2 scale, scaled by row. D. Heatmap illustrating the conversion of the role of ERK in naïve vs primed pluripotent cell types. The colour scale represents the log2 fold change (LFC) for each indicated sample between 4h of ERK activation and 0h. While in ESCs and RSCs ERK downregulates most pluripotent genes, in EpiLCs pluripotent genes are being upregulated upon ERK. E. The genes annotated as the GO term MAPK cascade (GO:0000165) were plotted in average expression changes (scaled from −1 to 1) between the conditions described in Fig 2A. Each line represents a gene. The thickest line represents the average of all genes. F. Barplots indicating the log2 fold change (LFC) of either 2h or 4h of ERK induction compared to the baseline at 0h, as indicated. Early activated ERK genes are upregulated at 2h in all cell types, while lineage specific late genes become up- or downregulated at 4h, showing cell type-specific behaviour.

Naïve ESC populations express naïve pluripotency markers, while EpiLC populations express primed pluripotency markers (Fig. 2C). RSCs, as an intermediate population, show a transcriptome that clusters between ESCs and EpiLCs (Fig. 2B) and express both naïve and primed markers (Fig. 2C). As expected ERK repressed pluripotency genes in naïve ESC (Hamilton et al., 2019), but in EpiLCs it induced them (Fig. 2D), consistent with the role of FGF in supporting primed pluripotency (Greber et al., 2010; Tesar et al., 2007). This included genes normally associated with naïve pluripotency, such as *Klf2*, *Klf4*, *Esrrb* and *Prdm14* (Fig. 2D). We confirmed that ERK-regulated genes defined in Serum/LIF ESCs (Hamilton et al., 2019) were also being up- and down-regulated in our ESC samples (Fig. S2A). Finally, we investigated whether we could detect differences between early (upregulated at 2h) vs late (at 4h) genes. Overall, the ERK transcriptional response could be separated between early ERK-responsive or MAPK canonical genes, that are upregulated in all cells (Figs 2E-F), and late responsive genes which were more specific to each cell type (Fig. 2F).

### ERK-mediated gene repression is cell type-specific while activation is more ubiquitous

Since we obtained a time resolved dataset, we defined genes regulated by ERK in all conditions by clustering the differentially expressed genes according to the dynamics of activation and repression in response to induction and inhibition of the pathway across time (Figs S3A-C, Table S1). For activated genes, we selected clusters with highest expression at 4h (cluster 2 ESC, cluster 3 RSC, cluster 3 EpiLC in Figs S3A-C). For repressed genes, we selected clusters where their lowest expression is at 4h (clusters 1 & 3 ESC, clusters 1, 2 & 4 RSC, cluster 4 EpiLC in Figs S3A-C). These clusters were compared between cell types to determine the extent to which ERK regulates common sets of genes in different contexts (Figs 3A-B, Table S2). While the strict overlap between cell type-specific activated genes was surprisingly low (Fig. 3A), the overall tendency trend of change is similar in all three cell types, with 70% of genes activated by ERK in ESCs exhibiting a positive fold change in EpiLCs, and the mean is always above zero (Fig. 3C). This contrasts with ERK-repressed genes, where almost no overlap is observed (Fig. 3B),only 50% of genes repressed in ESCs show negative fold changes in EpiLCs and the mean of the fold change is zero (Fig. 3D). Thus, while ERK-activated genes showed modest overlap between cell types, ERK-repressed genes were even more context-dependent (Figs 3A-B). For both repression and activation, the response in RSCs and naïve ESCs is far more similar than that observed in EpiLCs.

**Figure 3.**
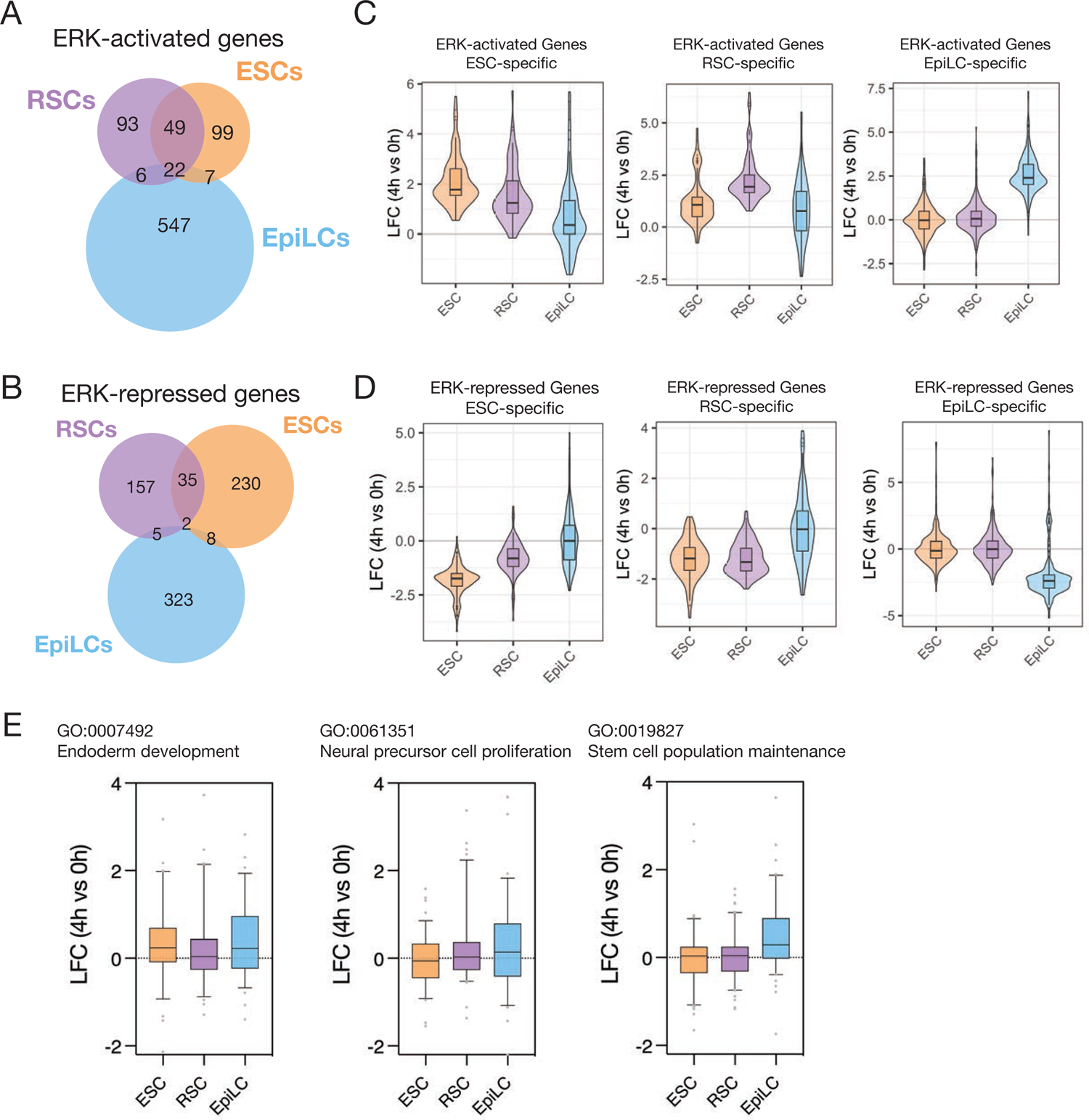
Repressed ERK targets are more specific to each cellular context than activated responses. A. Venn diagram showing the proportion of ERK-activated genes that are shared between cell types. The ERK-activated genes in each stage were defined from k-means clustering (See Fig. S3A-C and Methods) of all differentially expressed genes between all conditions (differentially expressed were defined as padj < 0.05 and abs (LFC) > 1.5). B. Venn diagram showing the proportion of genes repressed by ERK that are shared between cell types. The ERK-repressed genes in each stage were defined from k-means clustering (See Fig. S3A-C and Methods) of all differentially expressed genes between all conditions (differentially expressed were defined as padj < 0.05 and abs (LFC) > 1.5). C. Violin Plot of the log2 fold change (LFC) for each indicated cell type between 4h of ERK activation and 0h, for the activated set of genes found specifically in ESC, RSC and EpiLC, respectively. D. Violin Plot of the log2 fold change (LFC) for each indicated cell type between 4h of ERK activation and 0h, for the repressed set of genes found specifically in ESC, RSC and EpiLC, respectively. E. Violin Plot of the log2 fold change (LFC) for each indicated cell type between 4h of ERK activation and 0h, for the genes annotated as the indicated GO Term.

To understand the functional role of ERK-activated genes in each context, we extracted genes lists annotated as a specific GO Term and compared their fold changes upon ERK activation in each cell type. We found that genes annotated as endoderm related (GO:000792) were induced by ERK in both naïve and primed (Fig. 3E), consistent with the ability of these cell types to make primitive and definitive endoderm (Anderson et al. 2017). However, genes related to the neural lineage (GO:0061351) and stem cell maintenance (GO:0019827) were better induced by ERK in primed cells (Fig. 3E).

Given that ERK-repressed genes were more context-dependent than ERK-activated genes (Figs 3A-B), this suggests that repression of a set of genes might play a more important role in shaping specific responses than activation. We found genes related to placenta development repressed in RSCs and angiogenesis in EpiLCs (Fig. S3D, Table S3). These data support the idea that ERK activates genes related to lineage progression but in specific cell types it also has an important role in suppressing non lineage-specific gene expression.

### *In vivo* ERK responses recapitulate the *in vitro* transcriptome

To complement the analysis of *in vitro* responses to ERK in naïve vs primed stages, we sequenced the transcriptome from *in vivo* pre- and post-implantation Epiblast before and after ERK activation (Fig. 4A). PCA of the uninduced samples suggests that the major alterations in gene expression were due to *in vitro* adaptation, but it also illustrated a very clear separation of the naïve (ESCs and the pre-implantation Epiblast) from the primed (EpiLCs and the post-implantation Epiblast) states (Fig. 4B). To investigate the differences between *in vivo* and *in vitro*, we extracted the most significant genes driving variation across PC1, but we found them to be mostly related to biosynthesis and metabolism, as one might expect from *in vitro* adaptation, and not related to developmental processes (Fig. S4A). Moreover, E3.5 Epiblast cells expressed naïve pluripotency markers (Fig. 4C), similar to ESCs (Fig. 2C) while E5.5 Epiblast cells expressed primed pluripotency markers (Fig. 4C), similar to EpiLCs (Fig. 2C).

**Figure 4.**
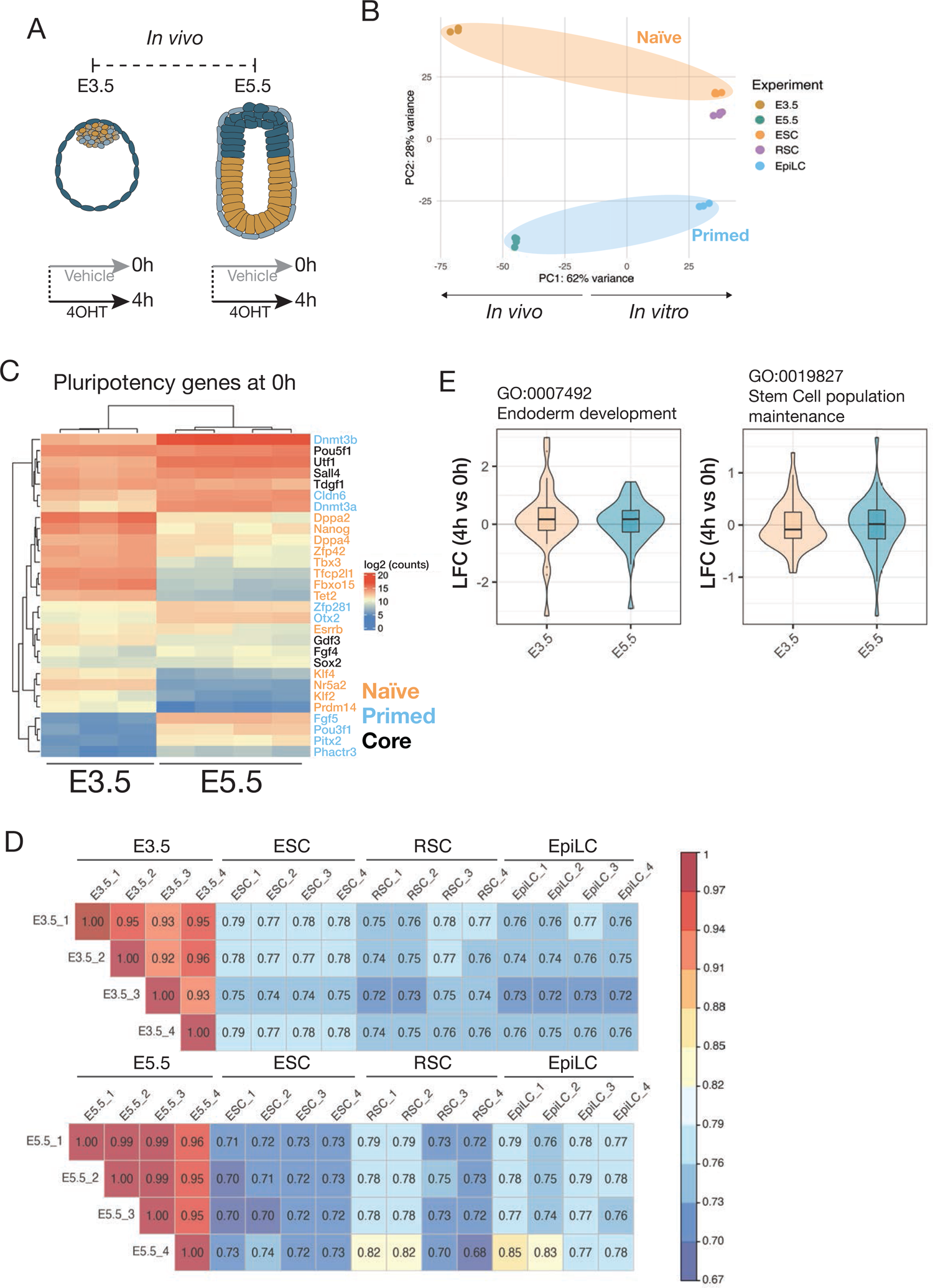
Overlap between *in vivo* and *in vitro* responses to ERK. A. Diagram of the *in vivo* populations collected for RNA-seq. Both pre-implantation (E3.5) and post-implantation (E5.5) Epiblast were isolated after 4h of treatment with either 4OHT (for ERK activation), or ethanol (Vehicle control). 4 replicates for each condition were collected, and 1 sample of E3.5 0h was discarded during Quality Control. B. Principal Component Analysis (PCA) of the 0h samples collected for RNA-seq shows that, while *in vitro* and *in vivo* populations cluster separately in PC1, PC2 separates naïve populations (ESCs and E3.5 Epiblast) from the primed populations (EpiLCs and E5.5 Epiblast), with the intermediate RSCs population in the middle. Each dot represents an individual biological replicate. ESCs n=4, RSCs n=4, EpiLCs n=3, E3.5 n=3, E5.5 n=4. C. Heatmap showing the expression of pluripotency markers (orange for naïve markers, blue for primed and black for shared or core pluripotency) of the *in vivo* populations before ERK induction. E3.5 Epiblast expresses higher expression of naïve markers than post-implantation E5.5 Epiblast, which expresses primed markers. Each column represents an individual biological replicate. E3.5 n=3, E5.5 n=4. The colour scale indicates normalised counts in log2 scale. D. Correlation matrix for *in vivo* vs *in vitro* 4h induced samples, showing higher correlation for E3.5 with ESC, and E5.5 with EpiLCs. Count matrices from RNA-seq were filtered for genes differentially expressed in E3.5 or E5.5, respectively, and covariances between all samples were calculated. Differentially expressed genes were defined as p_adj_ < 0.05 and abs (LFC) > 0.75). The colour scale indicates correlation coefficient. E. Violin Plot of the log2 fold change (LFC) for each indicated sample between 4h of ERK activation and 0h, for the genes annotated as the indicated GO Term.

We then computed correlation coefficients between *in vivo* and *in vitro* ERK-induced samples and found that ERK-induced E3.5 Epiblast correlated more closely to the ERK-induced ESC samples, while E5.5 Epiblast correlated better with EpiLCs (Fig. 4D). We also observed that the E3.5 Epiblast upregulated 58% of endoderm-related genes upon ERK induction (Fig. 4E), and E5.5 Epiblast showed 54% of stem cell-annotated genes being upregulated upon ERK induction (Fig. 4E), which recapitulated the effect observed *in vitro*.

While we see ERK inducing a global endoderm signature in E3.5 Epiblast (Fig. 4E), we saw more robust *Sox17* induction compared to *Gata4* or *Gata6* after 4h of ERK (Figs. 5A, S5A-C, Table S4). We therefore sought to validate the capacity of intracellular ERK induction to functionally promote GATA6-positive PrE in the blastocyst. We isolated E3.5 CRAFR26 embryos and treated them with either 4OHT or FGF4 for 4h. Then, the embryos were left to develop until late blastocyst stage (E4.5) in the presence of an FGFRi to prevent paracrine signalling (Fig. 5B). In ERK-induced blastocysts, we observed that 96% of the ICM cells expressed the PrE marker GATA6 at E4.5 (Figs 5B-C), and this induction was more efficient than the same length of treatment (4h) with FGF4 (Figs 5B-C). These surprising observations suggest that *Sox17* is induced independently by ERK, rather than downstream of GATA4 and GATA6, and that it may be one of the earliest immediate responses to ERK. Although GATA6 is believed to be an early marker of mouse PrE at a protein level (Artus et al., 2011), the induction of *Sox17* mRNA may be an earlier response to ERK than *Gata6* mRNA.

**Figure 5.**
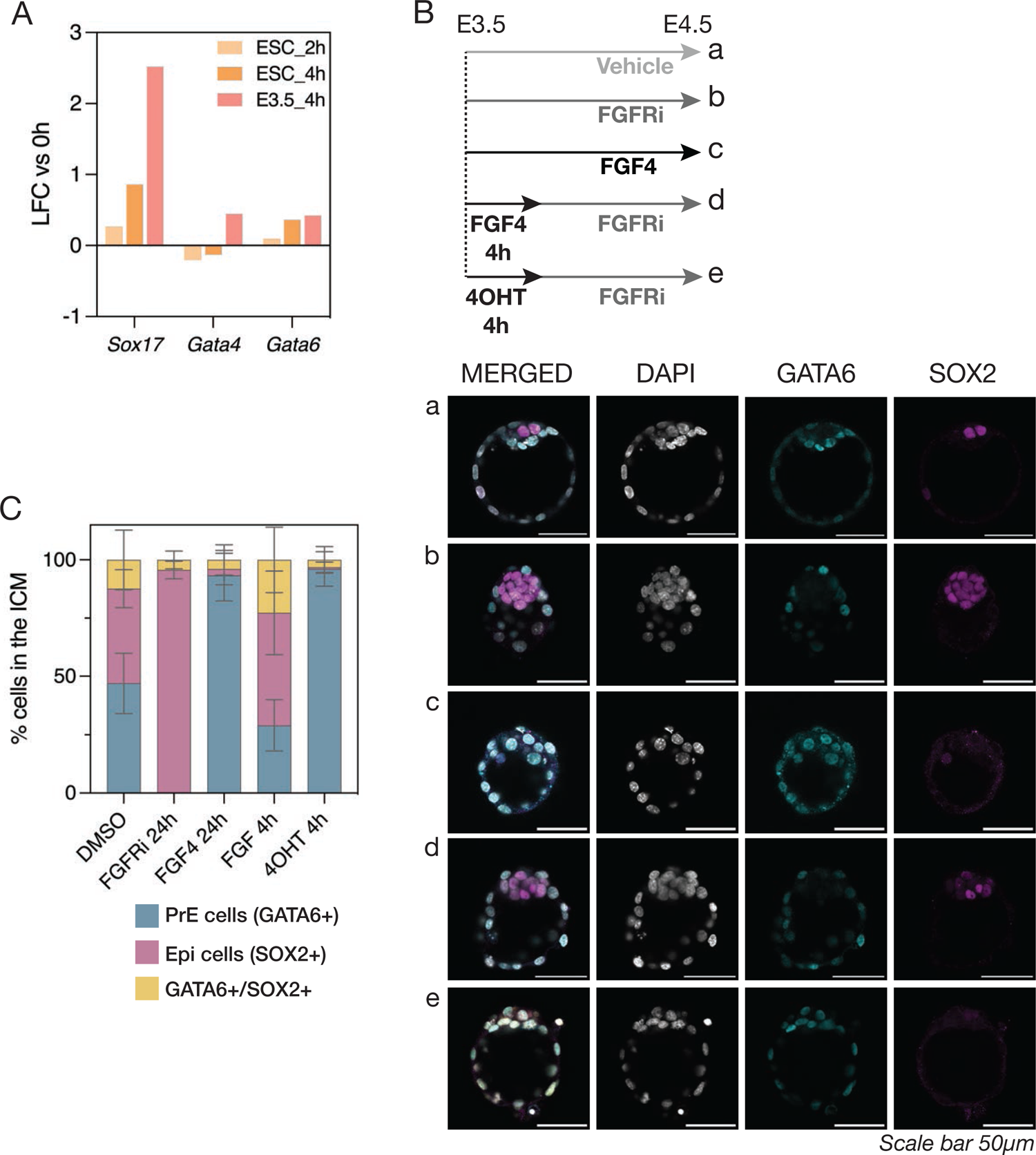
ERK induction in pre-implantation embryos is more efficient than FGF4 in inducing PrE. A. Barplot indicating the log2 fold change (LFC) of either 2h or 4h of ERK induction compared to 0h, as indicated. *Sox17* shows stronger transcriptional induction upon ERK than *Gata4* and *Gata6*, both *in vivo* (E3.5) and *in vitro* (ESC). B. E3.5 embryos were isolated and treated as indicated for 24h before fixing. Staining for SOX2 (Epiblast) and GATA6 (PrE) shows that ERK induction for 4h induces PrE at the same efficiency as FGF4 treatment for 24h. FGFRi (PD17) was added at 250 nM. FGF4 supplemented with heparin was added at 500 ng/ml. 4OHT was added at 250 nM. C. Quantification of immunofluorescence images from Fig. 5B. Only ICM cells were scored. PrE cells are scored at GATA6+ cells. Epiblast (Epi) cells are scored at SOX2+ cells. Double positive GATA6/SOX2 were also observed. None double negative cells were observed in the ICM.

### ERK triggers the remodelling of the chromatin landscape

We hypothesised that cell type-specific ERK responses could be mediated either by regulatory regions already differentially available when ERK signalling is activated, or by ERK triggering the reorganisation of the accessible regions to direct transcriptional changes. To distinguish between these two possible scenarios, we performed Assay for Transposase-Accessible Chromatin using sequencing (ATAC-seq) (Buenrostro et al., 2013; Grandi et al., 2022) (Fig. 6A), and assessed which regions were differentially accessible between cellular states (Table S5), and upon ERK induction (Table S6).

**Figure 6.**
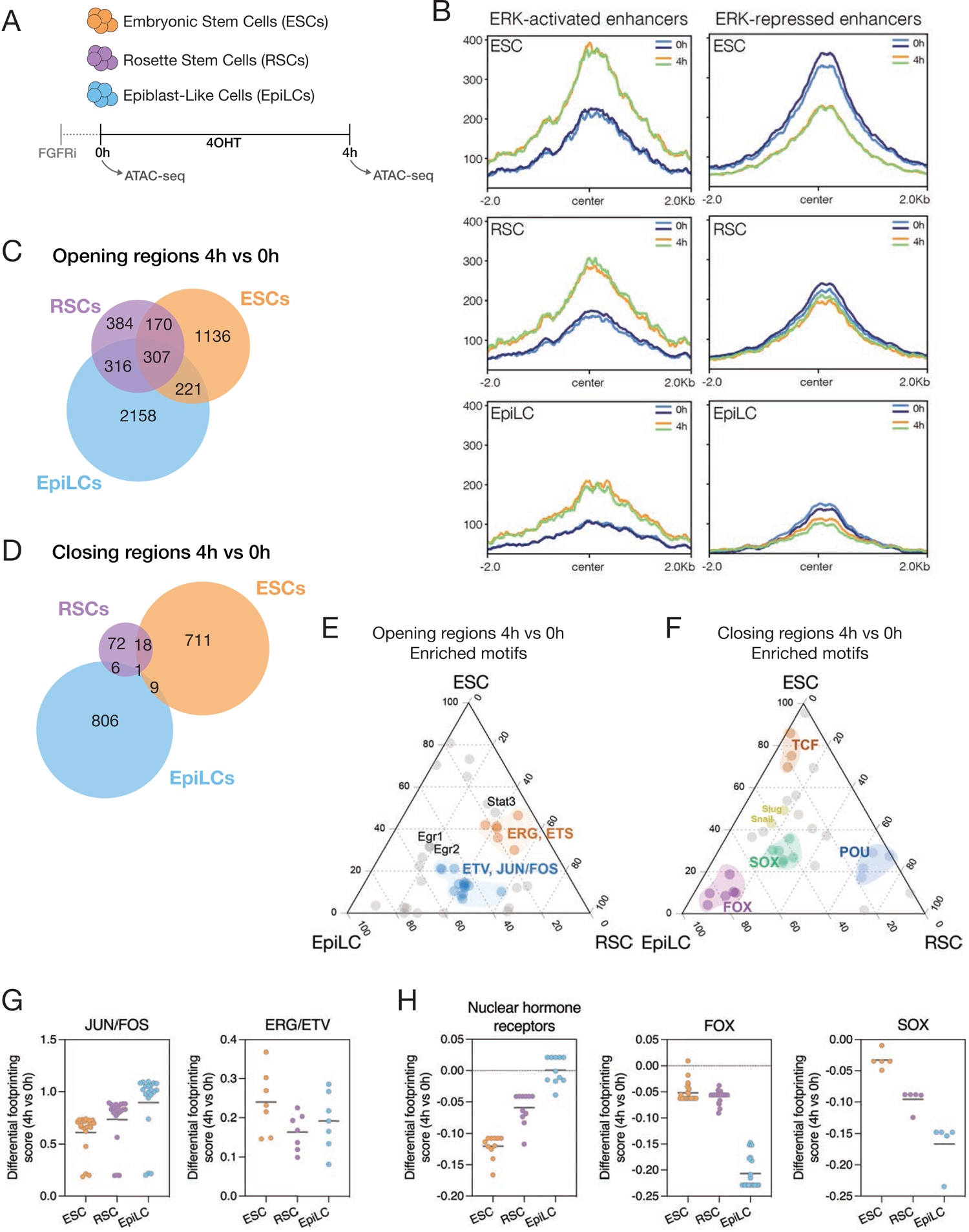
Chromatin accessibility cell-specific changes upon ERK induction *in vitro*. A. Diagram depicting the *in vitro* cell types collected for ATAC-seq, before and after 4h of ERK activation by addition of 4OHT. 2 biological replicates were collected for each condition. All cells were pre-treated with FGFRi for 48h before ERK induction, except for EpiLCs which were pre-treated for 4h. B. Scaled signal intensity from ATAC-seq of ESC, RSC and EpiLC comparing 0h and 4h of ERK induction, on the ERK-regulated enhancers defined in Hamilton et al., 2019. C. Venn diagram showing the overlap between regions that gained accessibility upon ERK induction (Opening regions, p_adj_ < 0.01 and LFC > 1.5), comparing between the different cell types described in Fig. 6A. D. Venn diagram showing the overlap between regions that lost accessibility upon ERK (Closing regions, p_adj_ < 0.01 and LFC < 1.5), comparing between the different cell types described in Fig. 6A. E. Ternary plot depicting the motifs enriched in the opening regions between 0h and 4h of ERK, comparing p-value scores across ESC, RSC and EpiLC. The axes represent -log10 of p-value. F. Ternary plot depicting the motifs enriched in the closing regions between 0h and 4h of ERK, comparing p-value scores across ESC, RSC and EpiLC. The axes represent -log10 of p-value. G. Footprinting predicted binding score were calculated for all consensus peaks at 0h and 4h. Binding scores are normalised to number of binding sites and accessible regions. Then a differential score is computed to illustrate the fold change in binding between 4h and 0h of ERK induction. TFs of the indicated families were grouped together, showing enriched binding of TFs from the canonical ERK factors (AP-1/ETS) upon ERK induction. H. Footprinting predicted binding score were calculated for all consensus peaks at 0h and 4h. Binding scores are normalised to number of binding sites and accessible regions. Then a differential score is computed to illustrate the fold change in binding between 4h and 0h of ERK induction. TFs of the indicated families were grouped together, showing distinct families of TFs being unbound upon ERK in specific cellular contexts.

We validated the changes observed by ATAC-seq in ESCs by exploiting our existing data on enhancer activity in response to ERK (Hamilton et al., 2019). We found that opening regions gained cofactor binding, including EP300 and Mediator (Med24) occupancy, as well as an increase in the active enhancer histone modification H3K27 acetylation (H3K27ac) (Fig. S6A). We also observed that regions that were becoming less accessible in response to ERK were losing EP300, Med24 or H3K27ac (Fig. S6A). Super enhancers, large cooperative DNA elements known to regulate transcription in pluripotent cells (Hnisz et al. 2013), were not remodelled upon ERK activation (Fig. S6B), consistent with our previous observations (Hamilton et al., 2019).

To validate cell type-specific changes in our dataset, we analysed the ATAC-seq signal at ESCs- and EpiLCs-specific enhancers (Buecker et al. 2014). We found that ESC-specific enhancers were more accessible in ESCs and closed down progressively in RSCs and EpiLCs, while EpiLC enhancers were more accessible in EpiLCs than in ESCs. (Fig. S6C). Interestingly, both sets of enhancers were at an intermediate accessible state in RSCs (Fig. S6C), although RSC expressed mostly naïve genes (Fig. 2C). Finally, we assessed changes at the set of ERK-activated and repressed enhancers we previously defined based on EP300 and H3K27ac changes (Hamilton et al. 2019.). We found that, although these enhancers were defined in naïve ESCs, ERK-activated enhancers appeared to become more accessible with signalling in all three cell types (Fig. 6B). In contrast, while ERK-repressed enhancers became less accessible in naïve ESCs, this phenomenon was negligible in RSCs and EpiLCs (Fig. 6B), providing further support for the notion that cell type-specific responses depend on signalling-mediated repression.

### ERK-activated regulatory regions are shared between cellular contexts while repressed regions are more cell type-specific

While we observed that the positively regulated naïve ESC enhancers were responding to signalling at some level in all three cell types, we wished to take a non-biased approach to identify cell type-specific ERK-regulated regions. Thus, we compared the sets of peaks that were opening and closing with ERK induction in each cell type. Consistent with the RNA-seq and the ERK-induced sequences (Fig. 6B), there is considerable overlap in regions that opening in response to ERK (Fig. 6C). The overlap of opening regulatory regions was considerable, with 307 shared regulatory regions between all three cell types (Fig. 6C), showing higher overlap than the induced gene sets (Fig 3A). ERK closed regions appear again almost mutually exclusive (Fig 6D), suggesting that the transcriptional mechanisms governing repression are more specific to each cell type, whereas activation is shared between cellular contexts.

To assess the commonalities in ERK-responsive regions, we extracted the enriched motifs in each set of opening regions and found that most motifs were shared between the 3 cell states and were related to ERK-canonical factors (ETS–AP-1 motifs) (Fig. 6E, Table S7). However, motifs in the closing regions were more enriched in specific contexts (Fig. 6F, Table S7). We found motifs related to naïve pluripotency being closed in ESCs and RSCs (TCF, OCT motifs), and motifs related to later development stages being decommissioned in EpiLCs (FOX, SOX motifs) (Fig. 6F, Table S7). To further confirm that repression appears cell type-specific while activation seems relatively ubiquitous, we determined the genes nearest to opening and closing regulatory regions. We found a large number of opening regions mapped to the same genes in different cell types, but a high degree of specificity in negative regulation (Fig. S6D).

We then wanted to discern whether the regulatory regions were merely being remodelled or were occupied by different Transcription Factors (TFs). We examined changes in TF occupancy based on the predicted capacity of a bound TF to block the activity of Tn5 transposase and create a footprint in the ATAC-seq signal. We exploited the computational tool TOBIAS (Bentsen et al., 2020) to extract TF footprints from ATAC-seq data and predict changes in TF occupancy upon ERK induction, obtaining a differential binding score. We found that the ERK-canonical TFs (ETS–AP-1) showed similar increased TF-binding scores upon ERK in all cells (Fig. 6G, Table S8). Consistent with this, we observed 53% of footprints to be shared in all cell types in the 307 commonly opening regions, with a core set of ETS–AP-1 TFs in 38% of these in all cell types (Fig. S6E, Table S8).

When we look for global TF binding release in response to ERK, we see little change in all three cell types (Fig. S6F). Notably, the small number of TFs that are significantly unbound show some cell type-specificity, with nuclear hormone receptors being unbound in ESCs and FOX/SOX TFs losing binding in EpiLCs (Fig. 6H). However, consistent with our previous observations, the great majority of TFs remain associated with binding sites in cis-regulatory elements following ERK-mediated repression, including pluripotency factors like SOX2 in naïve ESCs (Fig. S6F, Table S8). In summary, while ERK mediated activation appears to be triggered by canonical ETS–AP-1 TFs, cell type-specific repression appears to be mediated by decommissioning in the absence of robust reduction to lineage-specific TF binding.

### ERK mediates distinct cis-regulatory strategies in specific contexts

In response to ERK stimulation we observed alterations in accessibility of regulatory regions, both previously defined as enhancers and new regions reported here. We next sought to investigate whether this ERK-mediated remodelling had an impact on ERK-regulated genes. By combining the RNA-seq with the ATAC-seq, we could distinguish between two strategies: either ERK is regulating genes in the proximity of already accessible regions, or factors downstream of ERK are directly influencing accessibility to produce changes in gene expression.

A significant number of regions were specifically accessible in each cell type before ERK induction (at 0h) (Fig. S7A, Table S5), and these regions were enriched for different TF motifs (Fig. S7B, Table S7). When we analysed the Log2FoldChange (LFC) of the genes closest to these peaks, we found that in EpiLCs the genes found next to already accessible regions were more strongly upregulated than in ESCs and RSCs (Fig. 7A, Table S9). This suggested that signalling acts on accessible regulatory sequences in EpiLCs, as exemplified in the *Spred3* enhancer, progressively accessible in RSCs and EpiLCs, while its transcription is activated by ERK solely in EpiLCs (Fig. 7B).

**Figure 7.**
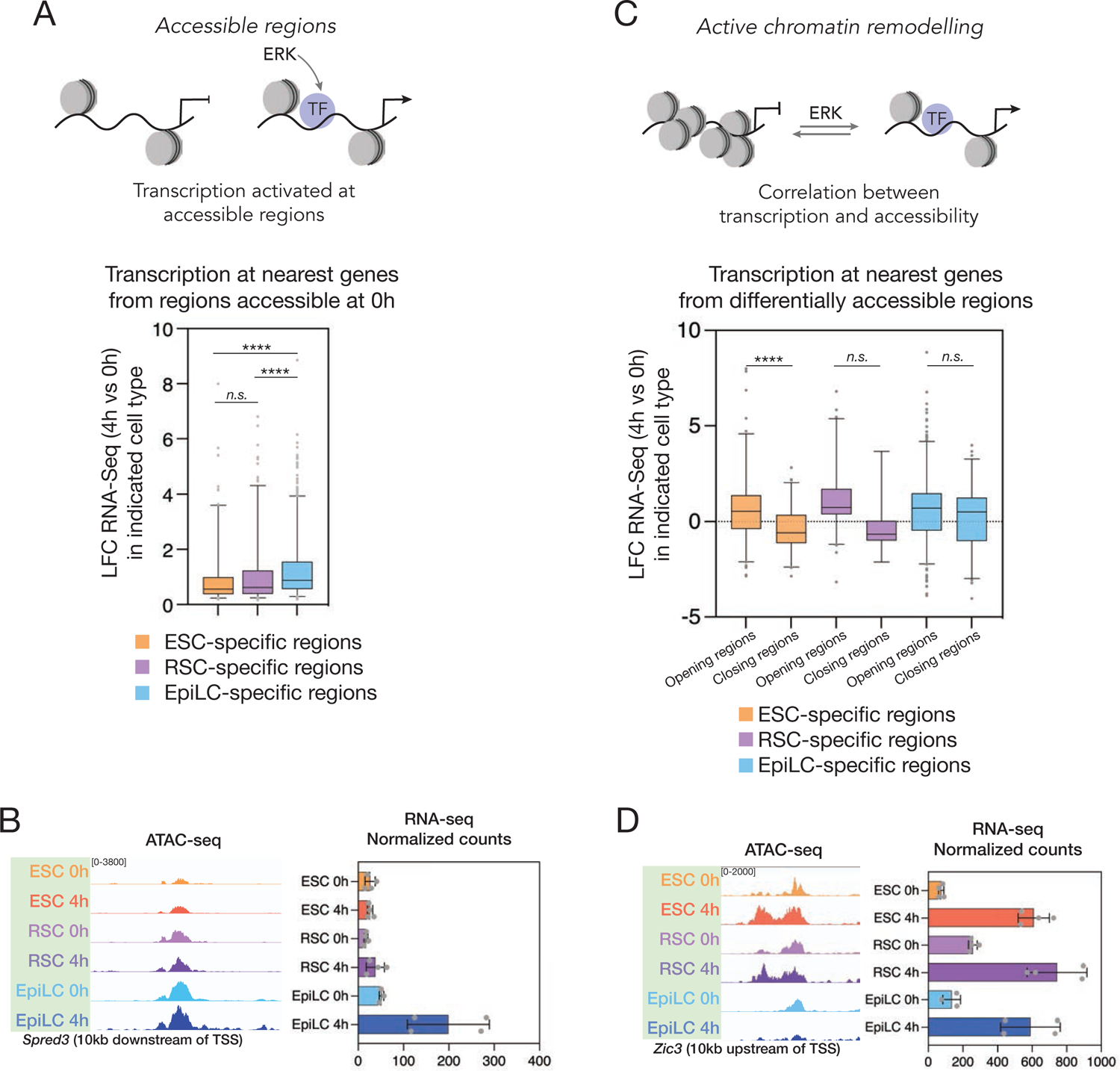
ERK mediates context-specific strategies in chromatin accessibility to drive transcriptional responses. A. The ATAC-seq peaks were classified as specifically accessible in each cell type at 0h as shown in Fig. S7A. Then, the log2 Fold Changes (LFC) of the transcription at the closest genes are shown for each category of peaks. RNA-seq LFC values were filtered for p_adj_ < 0.05. **** represents p-value < 0.0001 from Kruskal-Wallis test. n.s. represents a non-significant p-value > 0.01. B. Genome browser tracks (IGV v.2.16.0) showing ATAC-seq coverage for the *Spred3* enhancer and the corresponding normalized count for RNA-seq for the *Spred3* transcript. The regulatory region downstream of the TSS is available in RSCs and increasingly in EpiLCs before ERK induction (left) but transcription is only activated in EpiLCs upon ERK induction (right). C. The ATAC-seq peaks were classified as more (Opening) or less (Closing) accessible with ERK. Then, the log2 Fold Changes (LFC) of the transcription at the closest genes are shown for each category of regions. RNA-seq LFC values were filtered for p_adj_ < 0.05. **** represents p-value < 0.0001 from Kruskal-Wallis test. n.s. represents a non-significant p-value > 0.01. D. Genome browser tracks (IGV v.2.16.0) showing ATAC-seq coverage for the *Zic3* enhancer and the corresponding normalized count for RNA-seq for the *Zic3* transcript. We found similar induction of transcription in all cell types (right). However, the regulatory region downstream was being made accessible with ERK induction in ESCs and RSCs but not in EpiLCs (left).

The observation that available regions do not seem to be exploited by ERK in naïve ESCs suggested a requirement for remodelling in the ERK response in naïve ESC. We therefore investigated whether the regions remodelled by ERK were driving significant changes in gene expression, and found a strong correlation between changes in accessibility and transcription at the nearest gene in naïve ESCs, but not in the other two cell types (Fig. 7C, Table S9). This was measured as a significant different in transcriptional change from nearest genes to opening vs closing regions (Fig. 7C). This was manifested when we examined the *Zic3* enhancer. Its transcript was upregulated at a comparable fold change in all three cell types, while its enhancer was opening upon ERK induction in naïve cells but did not correlate with transcript levels in primed cells (Fig. 7D). Taken together our observations suggest that the chromatin landscape already present before the signal arrives can produce a significant impact in the signalling output, given that ERK exploits a preconfigured set of targets in EpiLCs, whereas in naïve ESCs it drives the rearrangement of its target loci.

## Discussion

How the same signal can produce different effects depending on the cellular or tissue context is a long-standing question in developmental biology. While FGF/ERK is a fundamental mediator of diverse processes, including proliferation, survival and differentiation (Dorey and Amaya, 2010; Turner and Grose, 2010), in numerous tissues across development (Corson et al., 2003), it is difficult to fathom how it achieves this diversity of regulation through a set of conserved immediate early TFs. To investigate the context-dependent cellular responses to ERK, we have developed a tool that works both *in vivo* and *in vitro* and allows for precise, homogeneous and controlled activation of ERK in the tissue and time of choice and found that context-dependent transcription appears to depend largely on the capacity of this pathway to supress cell type-specific transcription.

One of the advantages of our model is that it overcomes heterogeneity and feedback inhibition. Historically this has complicated the assessment of responses to signalling, as the immediate early response involves feedback inhibition. Our previous findings suggest that the range of enhancer regulation and phosphorylation induced intrinsically in naïve ESCs is the same as that induced by FGF4 in FGF4 mutant cells, but that ERK phosphorylation occurs synchronously throughout the culture. This is likely the reason that four hours of ERK induction *via* 4OHT converts the ICM to entirely GATA6-positive PrE over the course of a subsequent 24-hour period, similar to that achieved only when FGF4 is present for the full 24 hours. What does this suggest about the network involved in Epiblast-endoderm segregation, that has always been modelled based on GATA6-FGF-NANOG? Perhaps that the induction of GATA6 may not be a direct transcriptional response to FGF/ERK, but rather its levels are stabilised someplace downstream of the immediate early response that includes induction of *Sox17* mRNA.

Given the effective homogenous response achieved with the c-RAF model, we have been able to produce quantitative data on changes in enhancer and promoter accessibility that suggest cell type-independent regulatory regions that acquire similar signalling-induced accessibility in all cell types. However, regions that become decommissioned in response to signalling were found to be cell type specific.

In naïve ESCs, the regions losing accessibility upon ERK induction are characterized by the presence of TCF motifs (Fig. 6F). ESCs and RSCs are cultured in nearly identical conditions, with the only difference being that canonical WNT signalling is activated in ESC *via* GSK3 inhibition and inhibited in RSC with IWP2 (see Methods). In naïve ESCs, canonical ß-catenin signalling exploits TCF3 to support pluripotency, by associating with these regulatory regions and counteracting TCF3-mediated repression(Athanasouli et al., 2023; Watanabe and Dai, 2011). Yet canonical WNT signalling also induces PrE differentiation in ESC and can do so in virtually identical ESC media with one key difference, the removal of the FGF block (Anderson et al., 2017). How does the same pathway support both activities? In response to ERK activation in naïve ESCs, we found decommissioned regulatory regions enriched in TCF binding, suggesting an FGF-dependent alteration in WNT activity preventing it from activating the TCF dependent pluripotency network. As a result, WNT activation now triggers endoderm differentiation based on an ERK dependent set of accessible TCF targets.

In the Epiblast lineage, where WNT is inhibited prior to ERK signalling in RSCs, repression by ERK once again targets naïve ESC enhancers, but this time in regions enriched in POU/OCT motifs such that signalling promotes the transition into primed pluripotency (Fig. 6F). These regions could reflect the acquisition of bivalency at pluripotency enhancers in RSCs, enabling them to both move forward developmentally, while retaining the capacity to revert to naïve ESCs (Neagu et al., 2020). Finally in EpiLCs, where FGF/ERK supports self-renewal, we identify the FOX factors prominent in gastrulation stage differentiation.

Historically it has been difficult to reconcile the role of FGF/ERK in neural differentiation with so called default neural induction. We previously observed that anterior neural identity can be achieved in the presence of ERK inhibition (Hamilton and Brickman, 2014), and similar results were obtained when BMP receptor mutants were cultured in a block to the FGF receptor (Di-Gregorio et al., 2007). However, this contradicts experiments in the chick embryo (Linker and Stern, 2004; Lunn et al., 2007) and ESC differentiation (Kunath et al., 2007; Stavridis et al., 2007), where it was argued that naïve to primed transition requires ERK. Our findings provide a means to reconcile these observations, by suggesting that context-dependent repression by ERK drives endoderm differentiation in the presence of WNT signalling, but Epiblast differentiation in its absence. The presence of POU motifs in ERK repressed RSC enhancers could also suggest that FGF/ERK represses the overlapping pluripotent/neural programs regulated by POU and SOX proteins, such that signalling supports the maintenance of Epiblast identity over its differentiation towards anterior neural (Jaeger et al., 2011).

At a regulatory level, the notion that cell type-specific response is largely about repression has a certain logic. Immediate early genes activated downstream of ERK are widely studied and are likely to be context independent. This contrasts with repression, that depends on the regulatory program transcribed in the target cell type. While activation is more ubiquitous, relying on a conserved set of factors, it would not be entirely - as time and duration would eventually alter the transcriptional program in the responding cell type. Thus common immediate early TFs would be influenced by time and the emergent network downstream of the signal would interact with cell type specific factors to produce a unique response. For example, we observe pluripotency genes are induced in EpiLCs only after four hours, and are unlikely to be regulated directly by ETS–AP-1 factors.

While this work focuses on FGF/ERK, the question of context is universal to all signalling pathways. Here we show that these ubiquitous factors can act identically in different cell types, but their capacity to induce transcription is not always correlated with alterations in chromatin structure. We found that the cis-regulatory strategy by which a signalling generates a transcriptional output requires remodelling in naïve ESCs, but not in EpiLC, suggesting that the same signal can employ different strategies depending on the context. This signal adaptability to the cellular context and chromatin has also been observed in other settings during development (Delás et al., 2023), but the means by which these pathways can effect changes in remodelling in one cell type, but not require it in another remains unknown. However, it does suggest that the canonical view of signalling specificity being governed by chromatin context and accessibility is not correct, as in this instance we observe the same immediate early response, but in one instance this involves remodelling and other not, suggesting that the first wave of ERK induction can overcome chromatin accessibility.

Overall, our data provides a framework for signal-dependent context specificity where both transcriptional and chromatin changes in response to ERK are more universal in activation than repression. As we found that context-dependent repression is mediated mainly by deactivation of enhancers, leaving cell specific TFs bound to their targets, this response is, by definition, context-dependent. Activation of ERK-responsive genes relies largely on commonly activated genes triggered as immediate early response to ETS–AP-1 canonical factors, but this can be propagated in time through existing cell-specific states. Whether activation exploits chromatin remodelling or not would depend on the existence of prebound core TFs, which could render cells plastic to respond to signalling and differentiate in specific contexts (Hamilton et al., 2019; Knudsen et al., 2023; Redó-Riveiro et al., 2024).

## Materials and methods

### Generation of the CRAFR26 construct

The CRAFR26 construct was engineered from two previous versions of the BXB plasmid (Hamilton and Brickman, 2014; Hamilton et al., 2019) that were digested and ligated to obtain a BXB-T2A-mCherry plasmid. Then, this fragment was introduced downstream of a lox-stop-lox cassette using the BigT plasmid (Srinivas et al., 2001), and then between the ROSA26 locus homology arms using the R26 plasmid (Srinivas et al., 2001).

### Generation of CRAFR26 cells and mouse line

The CRAFR26 construct was transfected into mESCs using a published gRNA directed to the Rosa26 locus (Gu et al., 2018), and the Cas9 plasmid PX458 (Addgene, #48138). Clonal colonies were initially screened by PCR, and then further validated by Southern blot (see probe sequence in Table S10), Sanger sequencing and in house karyotyping. For *in vitro* ERK induction experiments, the cells were lipofected with a pCAG-CRE plasmid to remove the STOP codon. The cells were then validated by flow cytometry, western blot and immunostaining.

To generate the CRAFR26 mouse line, correctly targeted CRAFR26 mESCs (Agouti background) were injected into C57BL/6 mouse embryos and chimera contribution was assessed by fur colour. Male chimeras were crossed with C57BL/6 females to obtain heterozygous F1s. Pups were genotyped from ear snips by PCR.

Primers and genotyping PCR as follows:

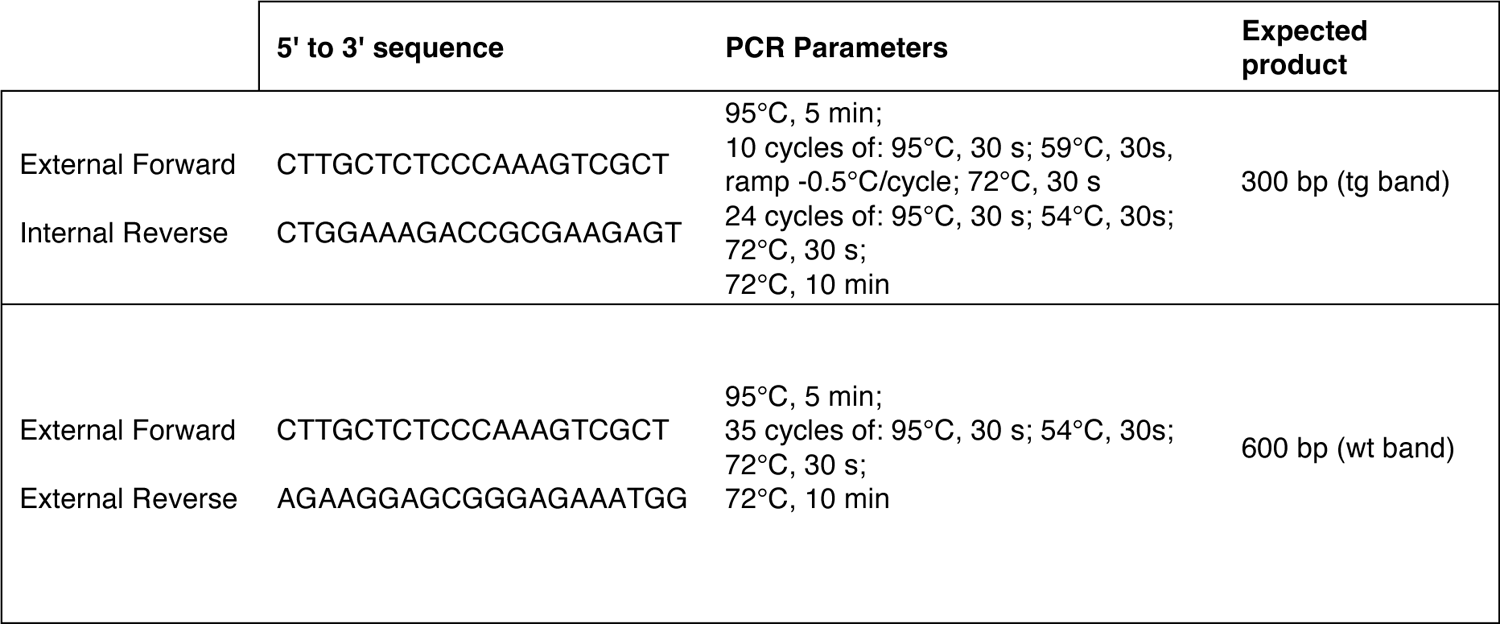

### Mouse husbandry

Mouse lines (*Mus musculus*) were maintained under 12h light cycles in designated facilities at the University of Copenhagen. Natural mating was set up in the evening, and females were checked for plugs the following morning, which was established as embryonic day 0.5 (E0.5). Mouse heterozygous Sox2Cre (Hayashi et al., 2002) females were mated with homozygous CRAFR26 males to produce constitutive CRAFR26 expression in pre-implantation embryos, and Sox2Cre males were mated with homozygous CRAFR26 females to produce expression of CRAFR26 in the Epiblast lineage in post-implantation embryos (Hayashi et al., 2003). Sox2Cre females were superovulated to obtain larger numbers of embryos per animal. Superovulation was performed by intraperitoneal injection of 5 IU pregnant mare serum gonadotropin (PMSG, Sigma) followed by 5 IU human chorionic gonadotrophin (hCG, Chorulon, Intervet) 42 to 50 hours later. The females were mated immediately following hCG injection.

Animal work was authorized by the Danish National Animal Experiments Inspectorate under project license 2018-15-0201-01520 and performed in accordance with Danish national guidelines and European legislation*mESCs culture* CRAFR26 mESCs were routinely cultured on 0.1% gelatine-coated flasks in 2i/LIF media (Ying et al., 2008) consisting of N2B27 media supplemented with 3 μM of the GSK3 inhibitor Chir (Axon, CHIR99021), 1 μM of the MEKi PD03 (PD0325901, Sigma PZ0162) and 1000 U/ml of LIF (prepared in house). N2B27 media was prepared as previously described (Knudsen et al., 2023). For passaging, confluent flasks were washed with PBS, incubated for 3 minutes in accutase (Sigma, A6964), and tapped to release the cells from the flask.

Accutase was inactivated by adding medium and the cells were centrifuged, resuspended and replated onto new gelatine-coated flasks. All mESCs lines used in this study were routinely karyotyped and tested for mycoplasma.

For differentiation into RSCs, mESCs were plated into N2B27 media supplemented with 2 µM IWP2 (Millipore, 681671), 1 µM of the MEKi PD03 (PD0325901, Sigma, PZ0162) and 1000 U/ml of LIF (prepared in house). After 3 days cells were passaged as RSCs (Neagu et al., 2020).

For EpiLC differentiation, RSCs were plated in clumps at high confluency (50,000 cells/cm^2^), on Fibronectin-coated plates (Millipore, FC010), into N2B27 media supplemented with 12 ng/ml bFGF (Peprotech, 450-33-100µG), 20 ng/ml Activin A (Peprotech, 120-14E-200µG) and 1 µM XAV939 (Sigma, X3004). Media was changed after 24h and EpiLCs were collected after 48h.

For ERK induction *in vitro*, mESCs and RSCs were cultured for 48h before the experiment in 250 nM of the FGFRi PD17 (PD173074, Sigma, P2499) instead of PD03. For EpiLCs, PD17 incubation was reduced to 2h prior to induction. Then, the c-RAF kinase construct was induced by adding 250 nM 4OHT (4-Hydroxytamoxifen, Sigma, H7904) to the media.

### Immunofluorescence staining of cells

Cells were cultured for immunostaining in 8-well slides (Ibidi, 80826), and then stained and imaged on the plate. Cells were fixed for 10 minutes with 4% Formaldehyde (Fisher Scientific, PI-28906) at room temperature, then washed 3 times with PBS, then treated with cold methanol for 10 minutes at −20°C and washed again 3 times with PBS. Cells were blocked for 2 hours at room temperature in 10% Donkey Serum (Sigma, D9663), 0.3% Triton X-100 (Sigma, T8787) in PBS. Primary antibodies (see Table S11) were incubated overnight at 4°C in 1% BSA (Sigma, A2153), then washed 3 times. Secondaries were incubated for 2 hours at room temperature also in 1% BSA, and further washed 3 times.

DAPI (Molecular Probes, D1306, 1 μg/ml) was then added for 5 minutes and washed twice with PBS. Cells were imaged in a Zeiss 780 confocal microscope.

### Isolation of mouse embryos

Timed pregnant females were euthanised by cervical dislocation. E2.5 embryos were flushed from the oviducts using M2 medium (Sigma, M7167), and E5.5-E6.5 embryos were dissected from the uterus. For Sox2Cre; CRAFR26 embryos, they were briefly imaged in a Deltavision Widefield Screening microscope to confirm their genotype by mCherry expression.

For longer culture of E2.5 embryos the zona pellucida was removed by brief incubation in acidic Tyrode’s solution (Sigma, T1788) and then the embryos were incubated in KSOM drops (Millipore, MR-101-D) covered in mineral oil (NidOil, Nidacon, NO-400K) at 37°C. To induce ERK, 4OHT (4-Hydroxytamoxifen, Sigma, H7904) was added at 250 nM for the times indicated. FGFRi (PD173074, Sigma, P2499) was added at 250 nM. FGF4 supplemented with heparin was added at 500 ng/ml. IVC2 media was used for culture of E5.5-E6.5 embryos (Bedzhov et al., 2014).

### Immunofluorescence staining of E4.5 mouse embryos

E4.5 embryos were fixed in 4% Formaldehyde (Fisher Scientific, PI-28906) at room temperature for 15 minutes, then washed twice in PBS. Embryos were permeabilized for 15 minutes in 0.25% Triton X-100 (Sigma, T8787) in PBS/PVP (3 mg/ml PVP (Polyvinylpyrrolidone, Sigma, PVP40) in PBS). Blocking was performed for 1 hour in 2% Donkey Serum (Sigma, D9663), 0.1% BSA (Sigma, A2153), 0.01% Tween20 (Sigma, P1379) in PBS/PVP. Primary antibodies (see Table S11) were added to blocking buffer and incubated overnight at 4°C. Embryos were washed 3 times for 10 minutes in blocking buffer, and then secondary antibodies were added to blocking buffer and incubated for 2 hours. Secondary antibodies were washed 3 times for 10 minutes in blocking buffer. DAPI (Molecular Probes, D1306, 1 μg/ml) was added to blocking buffer and incubated for 15 minutes at room temperature before washing once more in blocking buffer and mounted for imaging. Imaging was performed in a Leica Stellaris confocal microscope.

### Immunofluorescence staining of E5.5-E6.5 mouse embryos

E6.5 embryos were fixed in 4% Formaldehyde (Fisher Scientific, PI-28906) at room temperature for 30 minutes, and then washed 3 times for 10 minutes in PBS. Embryos were permeabilized for 30 minutes in 0.5% Triton X-100 (Sigma, T8787) in PBS. Blocking was done overnight in 2% Donkey Serum (Sigma, D9663), 0.1% BSA (Sigma, A2153), 0.1% Tween20 (Sigma, P1379) in PBS. Primary antibodies (see Table S11) were added to blocking buffer and incubated for 48h at 4°C. Embryos were washed 3 times for 15 minutes in blocking buffer, and then washed overnight. Secondary antibodies were added to blocking buffer and incubated overnight. Secondary antibodies were washed 3 times for 15 minutes and washed overnight in blocking buffer. DAPI (Molecular Probes, D1306, 1 μg/ml) was added to blocking buffer and incubated for 2 hours at room temperature before washing 3 times for 15 minutes in blocking buffer and mounted for imaging. All overnight incubations were done at 4°C with gentle shaking in the dark. Imaging was performed in a Leica Stellaris confocal microscope.

### Bulk RNA-sequencing (RNA-seq) of cells

RNA was extracted using the RNEasy Mini Kit (Qiagen, 74106) following manufacturer’s instructions. 4 biological replicates were collected per sample (ESCs vs RSCs vs EpiLCs) and per condition (0h vs 2h vs 4h vs rev), defined as 2 different passages of 2 independent clones. RNA was quantified by Nanodrop and 1 µg of RNA per sample was treated for ribosomal RNA depletion using the NEBNext rRNA Depletion Kit (NEB, E6350) and then immediately followed by Library prep using NEBNext Ultra II kit (NEB, E7770). Libraries were pooled and sequenced using a H75 kit from Illumina in a NextSeq500 sequencer following manufacturer’s instructions.

### Bulk RNA-sequencing (RNA-seq) of E3.5 mouse Epiblast

To obtain an ICM with only Epiblast cells, E2.5 Sox2Cre; CRAFR26 embryos were cultured in KSOM supplemented with 250 nM PD17 (PD173074, Sigma, P2499) for 24h. Then, to induce ERK, the media was supplemented with either 250 nM 4OHT (4-Hydroxytamoxifen, Sigma, H7904) for ERK induction, or ethanol at the same dilution for control conditions.

To collect the ICM, the trophectoderm layer was removed by blastocyst immunosurgery as previously described (Linneberg-Agerholm et al., 2023; Solter and Knowles, 1975). 4 biological replicates were collected per condition (treatment vs control), each containing a group of 10 ICMs. The samples were placed into a 1.5ml tube with lysis buffer from the Arcturus™ PicoPure™ RNA Isolation Kit (Applied Biosystems, KIT0204), and followed with RNA extraction according to manufacturer’s instructions. RNA was quantified using a Qubit™ RNA High Sensitivity (HS) kit (Invitrogen, Q32852). Libraries were prepared from 2 ng of RNA per sample using the NEBNext® Single Cell/Low Input RNA Library Prep Kit for Illumina (NEB, E6420). Libraries were pooled and sequenced using a P3-100 kit from Illumina in a NextSeq2000 sequencer following manufacturer’s instructions.

### Bulk RNA-sequencing (RNA-seq) of E5.5 mouse Epiblast

To induce ERK in E5.5 Sox2Cre; CRAFR26 embryos, they were cultured in IVC2 media (Bedzhov et al., 2014) supplemented with either 250 nM 4OHT (4-Hydroxytamoxifen, Sigma H7904) for ERK induction, or ethanol at the same dilution for control conditions. The Trophectoderm and Parietal Endoderm were removed by dissection using forceps. Then, the Epiblast was separated from the Visceral Endoderm by 8 minutes incubation on ice in a solution of 2.5% pancreatin (Sigma, P3292) and 0.5% trypsin (Sigma, T4799). 4 biological replicates were collected per condition (treatment vs control), each containing a group of 10 isolated Epiblasts. The samples were immediately placed into a 1.5 ml tube with 350 µl of lysis buffer, as per manufacturer’s instructions of the Micro RNEasy Kit (Qiagen, 74004), used for RNA extraction. After extraction, RNA was quantified using a Qubit™ RNA High Sensitivity (HS) kit (Invitrogen, Q32852). Libraries were prepared from 7 ng of RNA per sample using the NEBNext® Single Cell/Low Input RNA Library Prep Kit for Illumina (NEB, E6420). Libraries were pooled and sequenced using a P3-100 kit from Illumina in a NextSeq2000 sequencer following manufacturer’s instructions.

### Bulk RNA sequencing data analysis

Raw reads were processed with bcl2fastq (v 2.19.1) and STAR (v 2.5.3a) was used to map sequencing reads and to generate the count table (Dobin et al., 2013). Genes with less than 10 counts were discarded from downstream analysis. 1 sample of 0h EpiLCs and 1 sample of 0h E3.5 were discarded at Quality Control. Principal component analysis and differential expression analysis were performed using the DESeq2 (Love et al., 2014) and factoextra (Kassambara and Mundt, 2020) packages in R, significance was defined as abs(log2FC) > 1.5 and adjusted p < 0.05 for *in vitro* samples, and abs(log2FC) > 0.75 and adjusted p < 0.05 for *in vivo* samples. Principal component analysis was computed using the prcomp function. Hierarchical clustering was performed using the hclust function. Enrichment of GO terms was determined using the clusterProfiler package (Wu et al., 2021; Yu et al., 2012). Heatmaps were created using ComplexHeatmap (Gu et al., 2016).

### ATAC-seq

ATAC-seq or Omni-ATAC was performed as described in Grandi et al., 2022. Briefly, 50,000 cells were collected in a 1.5 ml DNA LoBind tube (Eppendorf, 0030108051), and lysed for 3 minutes on ice in 0.1% IGEPAL, 0.1% Tween-20, 0.01% Digitonin (Promega, G9441). Nuclei were washed and pelleted for 10 minutes at 500g, and then treated for 30 minutes at 37°C with Tn5 transposase (Illumina, 20034197). The DNA was extracted using DNA Clean and Concentrator 5-kit (Zymo Research, D4014), and libraries were prepared according to (Buenrostro et al., 2013). Libraries were quantified using the NEBNext Library Quant Kit (NEB, E7630S), then pooled at same molarity and paired-end sequenced with a P2-100 kit from Illumina in a NextSeq2000 sequencer following manufacturer’s instructions.

### ATAC-seq data analysis

Raw reads were processed with bcl2fastq, adapters were trimmed with cutadapt, reads were mapped to mm10 using bowtie2. ChrM, duplicate reads and reads in blacklisted regions were removed. Peaks were called using macs2 (v 2.2.7.1) (--broad -f BAMPE -- qvalue 0.001) and consensus peaks per condition were defined from two replicates per condition using the DiffBind package (Stark and Brown, 2011). We removed promoter regions using the publicly available Eukaryotic Promoter Database (Dreos et al., 2015; Périer et al., 1998). Reads within defined peaks were counted using subread featureCounts-p v2.0.3 (Liao et al., 2014). Counts were used to generate a count table for downstream differential enrichment analysis using DESeq2 (Love et al., 2014), to obtain the Log2 Fold Change (log2FC) enrichment per peak before and after ERK induction. Significance was defined as abs(log2FC) > 1.5 and padj < 0.01. Overlap of peaks between conditions was determined using bedtools intersect (v2.31.0) (Quinlan and Hall, 2010). Motif analysis and peak annotation was performed with HOMER (v4.11, findMotifsGenome.pl -size 200; and annotatePeaks.pl) (Heinz et al., 2010). Heatmaps and Profile Plots were created with deeptools (v3.5.1) (Ramírez et al., 2016). TF footprinting and differential binding score was determined using TOBIAS (Bentsen et al., 2020). Correction and scoring were done in merged bam files with both replicates, against all consensus peaks. BINDetect was performed in the peak set of interest. TF binding motifs for footprinting analysis were downloaded from the latest version of JASPAR (Rauluseviciute et al., 2023; Sandelin et al., 2004).

## Acknowledgements

We thank Heike Wollmann, Magali Michaut, Adrija Kalvisa and the reNEW Genomics Platform for technical expertise, support, and the use of instruments. We thank Jutta Bulkescher, Anup Shrestha and the reNEW Imaging Platform for training, technical expertise, support, and the use of microscopes. We thank Javier Martin Gonzalez, Ricardo Alonso Laguna Barraza and the Core Facility for Transgenic Mice for technical expertise and support. We thank the entire Brickman lab, especially Molly P. Lowndes and Rita S. Monteiro, for critical discussion and help with proofreading.

## Author contributions

MP and JMB conceived the study, designed and interpreted experiments. MP performed all experiments and analysed the data. MP and JMB wrote the manuscript.

## Data and materials availability

Previously published Hamilton et al. 2019 data that were used here are available in the Gene Expression Omnibus (GEO) repository under accession number GSE132444. Previously published Buecker et al. 2014 data that were used here are available in the Gene Expression Omnibus (GEO) repository under accession number GSE56138. The RNA-seq and ATAC-seq datasets generated in this study have been deposited in the Gene Expression Omnibus (GEO) repository and are available under these accession numbers: GSE259232 for RNA-seq in ESCs, GSE259233 for RNA-seq in RSCs, GSE259234 for RNA-seq in EpiLCs, GSE259235 for RNA-seq in E3.5, GSE259236 for RNA-seq in E5.5, GSE259237 for the ATAC-seq. Reagents generated in this study are available upon reasonable request to the corresponding author.

